# The adaptability of H9N2 avian influenza A virus to humans: A comparative docking simulation study

**DOI:** 10.1101/2020.05.21.108233

**Authors:** Hengyue Xu, Jiaqiang Qian, Yifan Song, Dengming Ming

## Abstract

Influenza A virus, the H9N2 subtype, is an avian influenza virus that has long been circulating in the worldwide poultry industry and is occasionally found to be transmissible to humans. Evidence from genomic analysis suggests that H9N2 provides the genes for the H5N1 and H7N9 subtypes, which have been found to infect mammals and pose a threat to human health. However, due to the lack of a structural model of the interaction between H9N2 and host cells, the mechanism of the extensive adaptability and strong transformation capacity of H9N2 is not fully understood. In this paper, we collected 40 representative H9N2 virus samples reported recently, mainly in China and neighboring countries, and investigated the interactions between H9N2 hemagglutinin and the mammalian receptor, the polysaccharide α-2,6-linked lactoseries tetrasaccharide c, at the atomic level using docking simulation tools. We categorized the mutations of studied H9N2 hemagglutinin according to their effects on ligand-binding interactions and the phylogenetic analysis. The calculations indicated that all the studied H9N2 viruses can establish a tight binding with LSTc although the mutations caused a variety of perturbations to the local conformation of the binding pocket. Our calculations suggested that a marginal equilibrium is established between the conservative ligand-receptor interaction and the conformational dynamics of the binding pocket, and it might be this equilibrium that allows the virus to accommodate mutations to adapt to a variety of environments. Our results provided a way to understand the adaptive mechanisms of H9N2 viruses, which may help predict its propensity to spread in mammals.

## INTRODUCTION

Avian influenza virus (AIV) H9N2 subtype is one of the most widely distributed subtypes of AIV in the world, causing huge economic losses to the poultry industry. It was first isolated at the turkey fork in the United States in 1966[1], and later reported in the Middle East, South Africa, Europe and the rest of the world[2,3]. In China, the H9N2 influenza virus was first found in chicken farms in 1992, which was found in the report of the outbreak of avian influenza in mainland China[4]. Since then, sporadic outbreaks of avian influenza, mainly H9N2, have occurred in Guangdong Province[5–8]. In 1998, a nationwide outbreak of H9N2 avian influenza occurred in China in several months, first in Hebei Province, and then rapidly spread to most chicken farms across the country[9]. Although H9N2 subtype has been proved to be a low virulent virus[10], it was shown that H9N2 is intrinsically related to H5N1, H7N9 and other highly pathogenic AIV and other epidemic strains[11]. In recent years, due to the threat of H9N2 subtype avian influenza virus to the safety of poultry industry and public health, more and more attention has been paid to the study of its infection and transmission mechanism.

Three-dimensional structure models provided valuable knowledge for the study of infection, adaptation and transmission mechanism of avian influenza virus. Infection of AIVs usually starts with the binding of the viral protein haemagglutinin A(HA) to the sialic acid receptor on the surfaces of the respiratory epithelial cells, causing membrane fusion and leading the virus to enter the host cell[12]. In birds, the receptors are mainly α-2,3-linked lactoseries tetrasaccharide a (LSTa) in *trans* conformation, while in mammals, they have *cis* conformation of α-2,6-linked lactoseries tetrasaccharide c (LSTc)[13]. Mutagenesis analyses highlighted conserved infection-initiating residues on the 190 helix (index based on H3 influenza virus’ sequence) [14]. In H9N2 subtype AIVs, HA protein has a strong tendency to bind to LSTa, but accumulated evidences show that they can also bind to LSTc under certain conditions, thus directly infecting mammals and humans[15,16]. *Particularly, the growth rate of L226-type H9N2 virus in human airway epithelial cells was found 100 times faster than that of Q226-type H9N2 virus, indicating L226 critically important for infecting mammals[17]*.

However, due to the high mutation rate in the binding pocket and the flexibility of binding glycan, most structural mode HA proteins binding with ligands are still lacking, which makes it difficult to evaluate the effect of virus mutations. In this paper, the structural models of 40 representative H9N2 strains circulated in China and neighboring countries last few years were analyzed, and the docking mode of binding pockets was studied and the mutation principle of virus infection and transmission was deduced and discussed.

## Material and methods

### Collection of HA protein sequence of H9N2 strains in China

In order to study the human infection and animal-to-human/human-to-human transmission of H9N2 subtype avian influenza virus in China, 40 strains of H9N2 influenza virus were collected, which often appeared in recent studies[10,18]. Three quarters of the selected strains were reported to have occurred in China from 1997 to 2012. The rest included three strains from South Korea, three strains from Japan, one strain from Israel, one strain from South Africa and two strains from the United States. Of all the known H9N2 strains, one of the two strains from the United States was the first one discovered in 1966. The selected strains are listed in Table S1.

### Structural homology modeling of selected HA1 proteins

In this work, the HA1 domain of studied proteins, which including the key 190- and 220-loop, is considered for structural homology modeling and docking analysis, taking swine H9N2 influenza virus hemagglutinin (PDB entry code 1JSI[13]) as reference model to bind the penta-saccharide LSTc (α-2,6). The selected HA1 proteins have a very similar sequence to the reference one, with an alignment score of 83 to 94 as determined using CLUSTAL2.0[19]. Structure homology modeling is implemented by software package Modeller version 9.4[20]. In each model, a water molecule was carefully preserved because it was proved to mediate the formation of hydrogen bonds between the sialic acid and Gly-228[13].

### Docking the LSTc to the binding pocket in H9N2 HA1 protein

We only used LSTc as ligand, and dock it with the above-motioned target protein HA using Autodock Vina (version 1.1.2)[21]), with a grid centered on atom C6 of the sialic acid of LSTc and a parameter file for docking simulation were also generated by AutoDockTool. A total of 18 rotatable bonds were assigned to LSTc, and the hydrogen atoms were fixed on their attached heavy atoms. The Lamarckian genetic algorithm was used to search LSTc confirmation after up to 200000 times of energy evaluation.

Considering the high structural similarity between the studied H9N2 proteins and the reference model, we expected LSTc also to have similar confirmation as in the X-ray crystal structure. Therefore, for a given HA1 protein, we selected an LSTc conformation from the docking decoy set, which has the smallest atomic root mean square deviation (RMSD) to the reference LSTc structure. Further, we assigned a weight functions of 1.0, 1.0, 0.4, 0.4 and 0.2 to the five subunits: the sialic acid moiety Sia-1, Gal-2, GlcNAc-3, Gal-4, Glc-5 respectively, where the smallest weight function was assigned to Glc-5 since it locates outside the binding pocket and have the least contribution of the ligand-binding. For each HA1 protein selected, two independent docking simulation calculations were carried out and compared.

### Characterizing the binding microenvironment in the framework of local atom geometry

In docking, contact distances are very important because they usually measure the overall strength of ligand binding affinity. Furthermore, other geometric quantities, such as relative contact orientations, can also be important in evaluating the ligand binding ability. For this sake, we constructed a local affine coordinate system, called this Leu226 coordinates, by using three atoms of Leu226 sidechain, namely CG, CD1 and CD2. The first two axes, and, represent the directions from CG to CD1 and that from CG to CD2, respectively. The third axis then take the direction of the cross product: ( A similar affine coordinate system, called Q226 coordinates, was also built by using the three heavy atoms of Q226, namely CD, OE1 and NE2, and gave three axes along the directions). These two local coordinates provide quantitative characterization of the contact configurations formed by the ligands, particularly the sialic acid Sia-1, with residue 226. Particularly, distribution of the heavy atoms of Sia-1, namely the Sia-1-O8, Sia-1-O1A, Sia-1-O1B, Sia-1-C1 were then analyzed and compared using these two local coordinates.

### Phylogenetic analysis of using key residual arrays

Sequence alignment analysis revealed that only half of the sequences of the H9N2 strains studied were completely conserved, and 34% of the remaining half are partially conserved. Among the mutations, we noted that only a few residues are actually interacting with LSTc either directly or indirectly through some intermediate water molecules. To assess the importance of these key mutations, for each virus, we made a much shorter sequence simply by sequentially combining amino acids at these sites. We then constructed a new phylogenetic tree with these short sequences by using MEGA5[22]. These phylogenetic thus represent the evolution process of the infection and transmission of H9N2 strain, as solely determined by its LSTc-binding feature. We then compared these trees with those derived using the whole-sequence alignment, thus measuring the importance of the key mutation sites.

## Results and Discussions

### Homologous structures of the H9N2 HA1 proteins are very conservative

The sequence similarity between the HA1 proteins of H9N2 strained studied and the swine influenza HA1 protein was evaluated by the program Clustal W 2.0[23]. The alignment scores were determined to be between 83 and 94, and the alignment score for the identical sequences was set to 100. The three-dimensional structure superposition calculations showed that the homologous structures generated by the MODELLER program[20] is very similar to the X-ray crystal structure (PDB code 1JSI[13]) (see Figure 1). The overall structure of the SIA-binding domain of the HA1 protein, i.e. the residues between Ala109 and Gly252 as in PDB 1JSI, was found to be very conservative (see Figure S1), and the averaged backbone atomic root-mean-square-derivation (RMSD) between the models and the X-ray crystal structure is 0.22 Å with a standard error of 0.03Å. In particular, the side chain atoms of residues (such as Trp143) that are closely bound to Sia-1 overlap well with the X-ray structure with the RMSDs smaller than 0.3Å. These results suggest that the overall 3D structures of H9N2 HA1 proteins, especially those portions bound to Sia-1, is very conservative, and that most mutations occur on residues with less exposure to Sia-1. Thus, it is reasonable to assume that the homologous modeling results in structures that bind to Sia-1 in a similar manner, as observed in the X-ray structure of the H9 swine HA1-LSTc complex.

**Figure 1.**
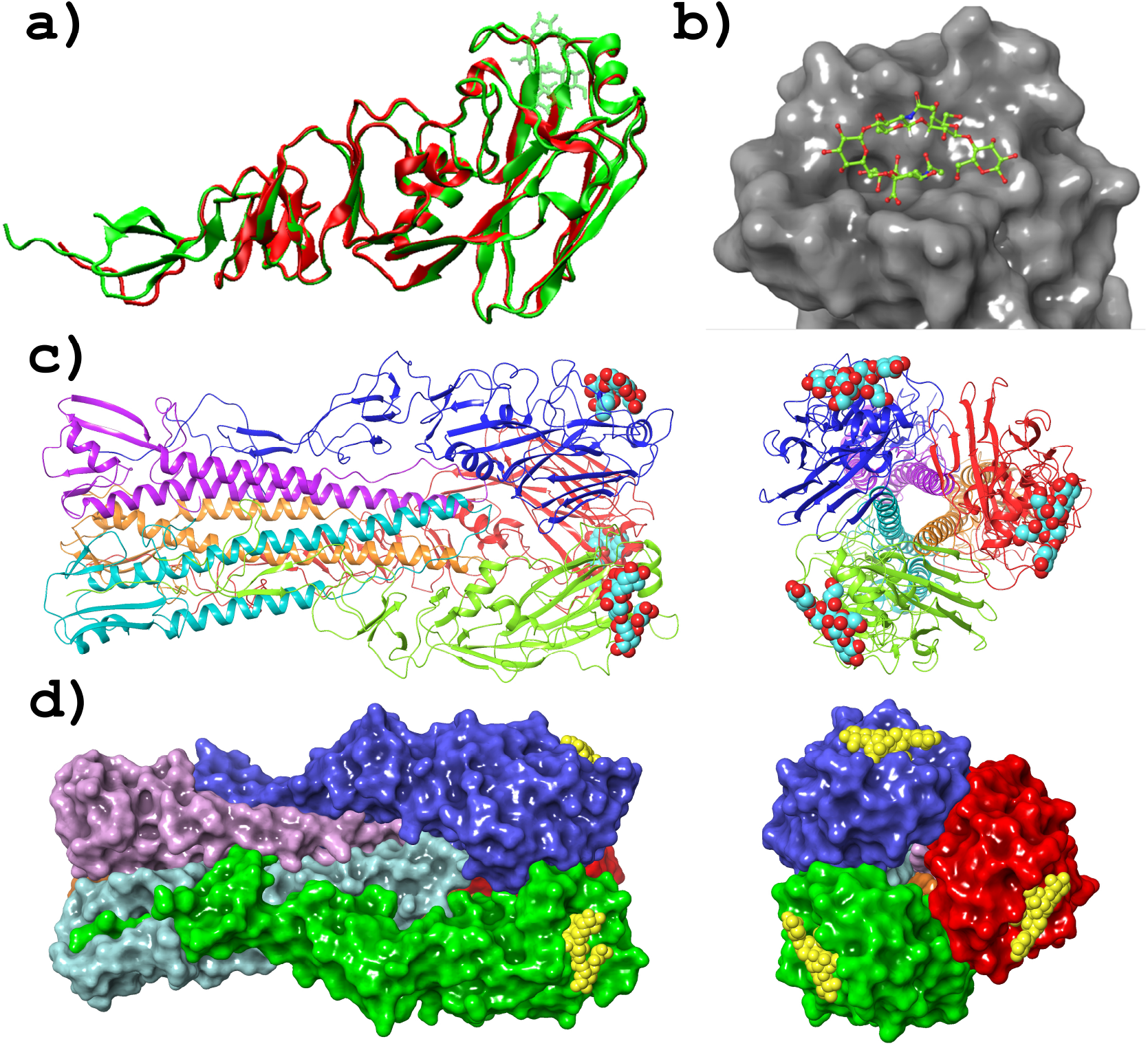
a) A typical homologous model (red) built by Modeller is very similar to the X-ray crystal structure (green, PDB code 1JSI), especially at the LSTc binding domain. b) the conservative binding pocket where LSTc is shown with ball-and-stick model. The structure of HA trimer is shown as cartoon c) and as surface d). This plot was prepared by PyMOL.

### Comparison of LSTc conformations in two dockings

We noticed that due to the large size of the ligand binding molecule LSTc and the fact that the binding pocket is not deep inside the protein but on the surface, the three units of the second half of the penta-saccharide ligand, namely GlcNAc-3, Gal-4 and Glc-5, fall on the outside of the binding pocket and are less constrained. A visual observation showed that the conformation of these three units predicted by docking varies greatly, while the calculated binding energy remains almost constant. Therefore, in order to determine the ligand-binding conformation in docking calculations, in addition to the low binding energy, we also require that the conformation of LSTc is the most similar to its conformation in the crystal structure. This requirement ensures the repeatability of the Vina docking outputs. Table 1 lists the results of the twice docking of LSTc with H9-HA1 proteins from studied strains. The results show that the LSTc conformations given by the two docking calculations are very close to each other. The following analyses on the ligand-binding features were then based on the complex structures of the second docking calculations.

**Table 1.**
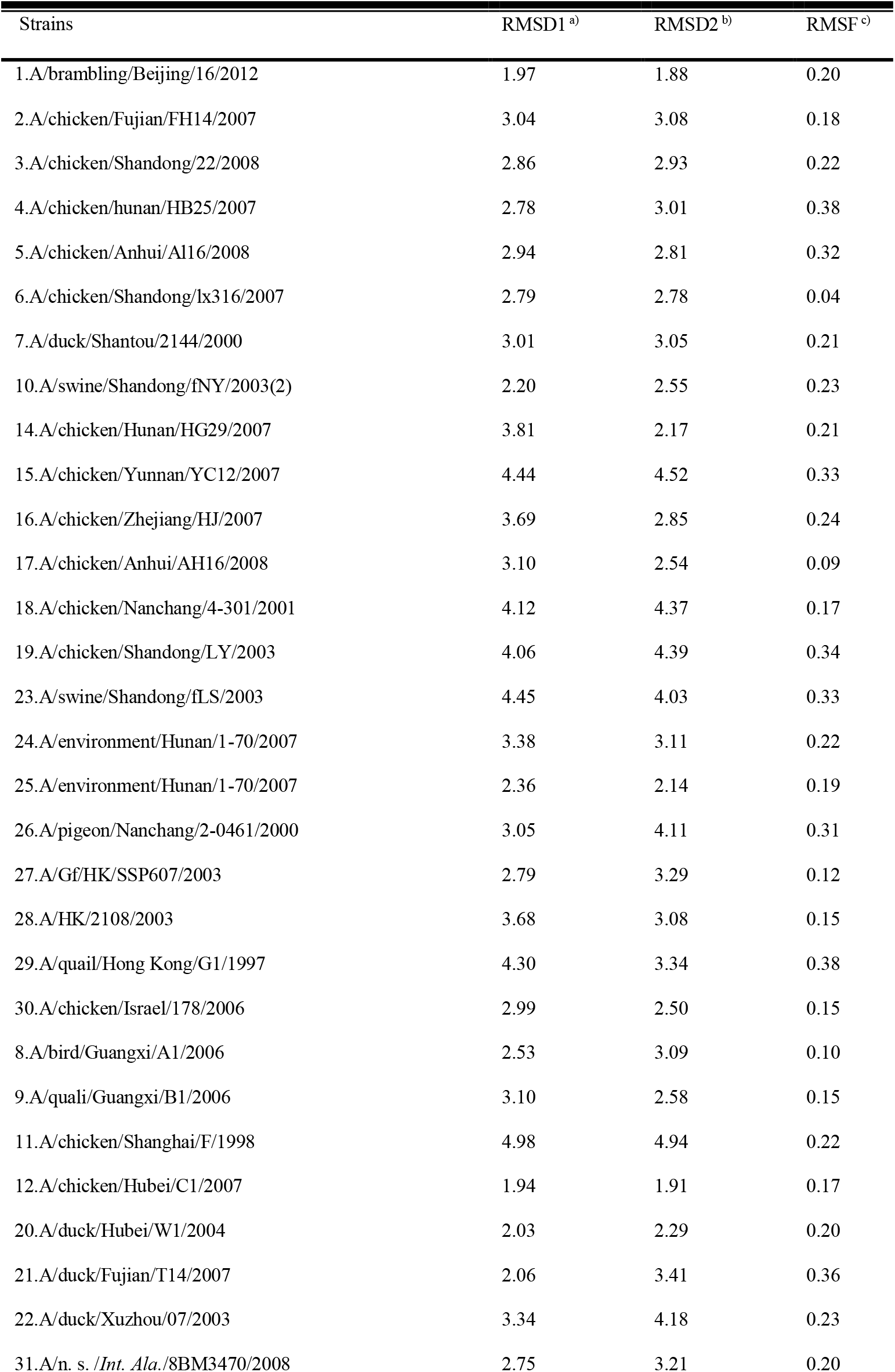
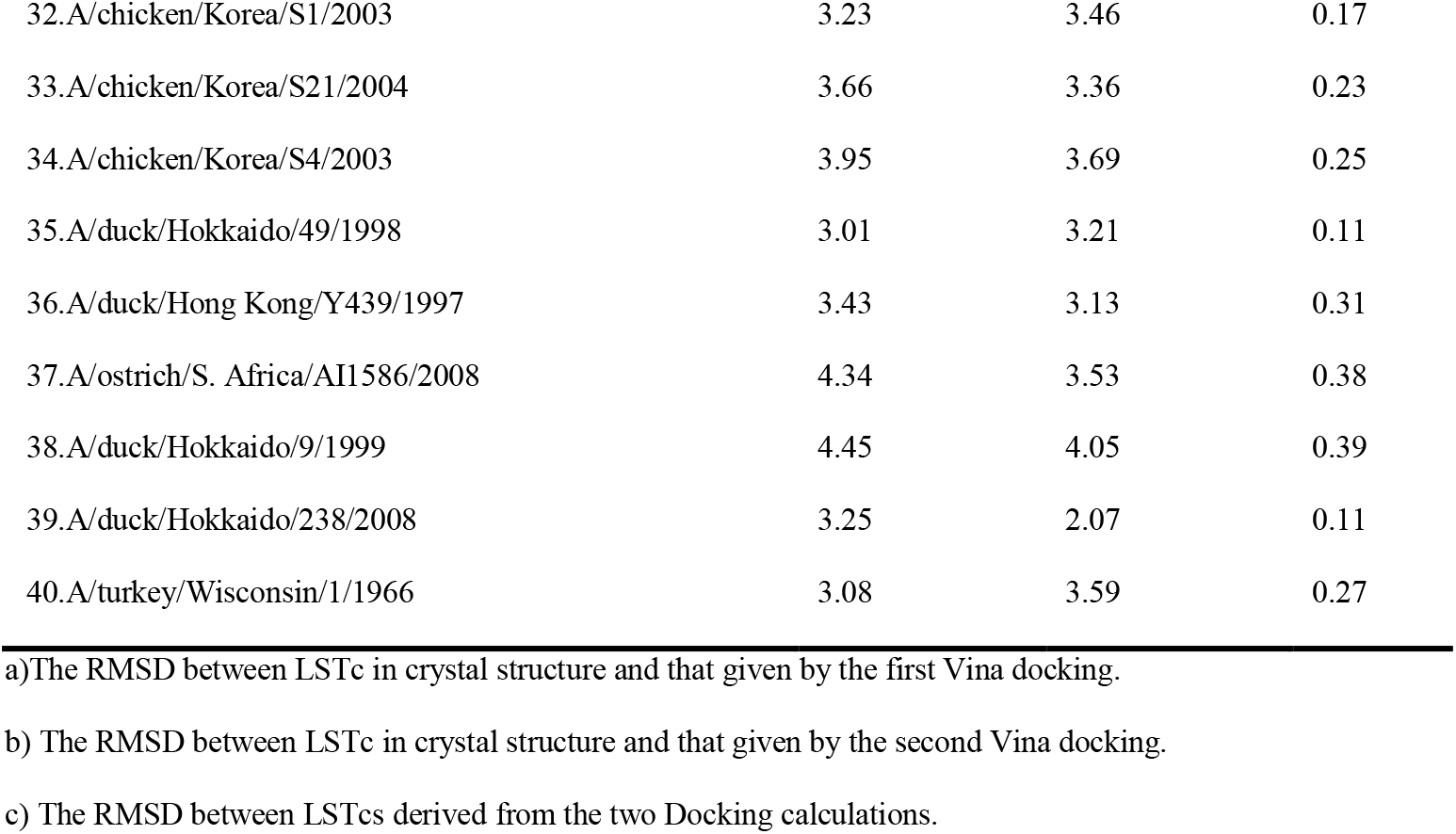
Comparison of the twice Vina docking

| Strains | RMSD1 <sup>a)</sup> | RMSD2 <sup>b)</sup> | RMSF <sup>c)</sup> |
| --- | --- | --- | --- |
| 1.A/brambling/Beijing/16/2012 | 1.97 | 1.88 | 0.20 |
| 2.A/chicken/Fujian/FH14/2007 | 3.04 | 3.08 | 0.18 |
| 3.A/chicken/Shandong/22/2008 | 2.86 | 2.93 | 0.22 |
| 4.A/chicken/hunan/HB25/2007 | 2.78 | 3.01 | 0.38 |
| 5.A/chicken/Anhui/A116/2008 | 2.94 | 2.81 | 0.32 |
| 6.A/chicken/Shandong/lx316/2007 | 2.79 | 2.78 | 0.04 |
| 7.A/duck/Shantou/2144/2000 | 3.01 | 3.05 | 0.21 |
| 10.A/swine/Shandong/fNY/2003(2) | 2.20 | 2.55 | 0.23 |
| 14.A/chicken/Hunan/HG29/2007 | 3.81 | 2.17 | 0.21 |
| 15.A/chicken/Yunnan/YC12/2007 | 4.44 | 4.52 | 0.33 |
| 16.A/chicken/Zhejiang/HJ/2007 | 3.69 | 2.85 | 0.24 |
| 17.A/chicken/Anhui/AH16/2008 | 3.10 | 2.54 | 0.09 |
| 18.A/chicken/Nanchang/4-301/2001 | 4.12 | 4.37 | 0.17 |
| 19.A/chicken/Shandong/LY/2003 | 4.06 | 4.39 | 0.34 |
| 23.A/swine/Shandong/fLS/2003 | 4.45 | 4.03 | 0.33 |
| 24.A/environment/Hunan/1-70/2007 | 3.38 | 3.11 | 0.22 |
| 25.A/environment/Hunan/1-70/2007 | 2.36 | 2.14 | 0.19 |
| 26.A/pigeon/Nanchang/2-0461/2000 | 3.05 | 4.11 | 0.31 |
| 27.A/Gf/HK/SSP607/2003 | 2.79 | 3.29 | 0.12 |
| 28.A/HK/2108/2003 | 3.68 | 3.08 | 0.15 |
| 29.A/quail/Hong Kong/G1/1997 | 4.30 | 3.34 | 0.38 |
| 30.A/chicken/Israel/178/2006 | 2.99 | 2.50 | 0.15 |
| 8.A/bird/Guangxi/A1/2006 | 2.53 | 3.09 | 0.10 |
| 9.A/quali/Guangxi/B1/2006 | 3.10 | 2.58 | 0.15 |
| 11.A/chicken/Shanghai/F/1998 | 4.98 | 4.94 | 0.22 |
| 12.A/chicken/Hubei/C1/2007 | 1.94 | 1.91 | 0.17 |
| 20.A/duck/Hubei/W1/2004 | 2.03 | 2.29 | 0.20 |
| 21.A/duck/Fujian/T14/2007 | 2.06 | 3.41 | 0.36 |
| 22.A/duck/Xuzhou/07/2003 | 3.34 | 4.18 | 0.23 |
| 31.A/n. s. /Int. Ala./8BM3470/2008 | 2.75 | 3.21 | 0.20 |
| 32.A/chicken/Korea/S1/2003 | 3.23 | 3.46 | 0.17 |
| 33.A/chicken/Korea/S21/2004 | 3.66 | 3.36 | 0.23 |
| 34.A/chicken/Korea/S4/2003 | 3.95 | 3.69 | 0.25 |
| 35.A/duck/Hokkaido/49/1998 | 3.01 | 3.21 | 0.11 |
| 36.A/duck/Hong Kong/Y439/1997 | 3.43 | 3.13 | 0.31 |
| 37.A/ostrich/S. Africa/AI1586/2008 | 4.34 | 3.53 | 0.38 |
| 38.A/duck/Hokkaido/9/1999 | 4.45 | 4.05 | 0.39 |
| 39.A/duck/Hokkaido/238/2008 | 3.25 | 2.07 | 0.11 |
| 40.A/turkey/Wisconsin/1/1966 | 3.08 | 3.59 | 0.27 |

a) The RMSD between LSTc in crystal structure and that given by the first Vina docking.
b) The RMSD between LSTc in crystal structure and that given by the second Vina docking.
c) The RMSD between LSTcs derived from the two Docking calculations.

### Mutations in the HA1 protein cause extensive perturbation of LSTc binding

Although the overall backbone structures of the H9 HA1 proteins studied are very conservative, we found that many of the mutations that occur in the binding pocket have a significant impact on the interaction between the protein and the sialic acid ligand. We categorized the strains studied into four groups based on the type and location of the mutations and their effects on protein-ligand interactions (Table 2).

**Table 2.**
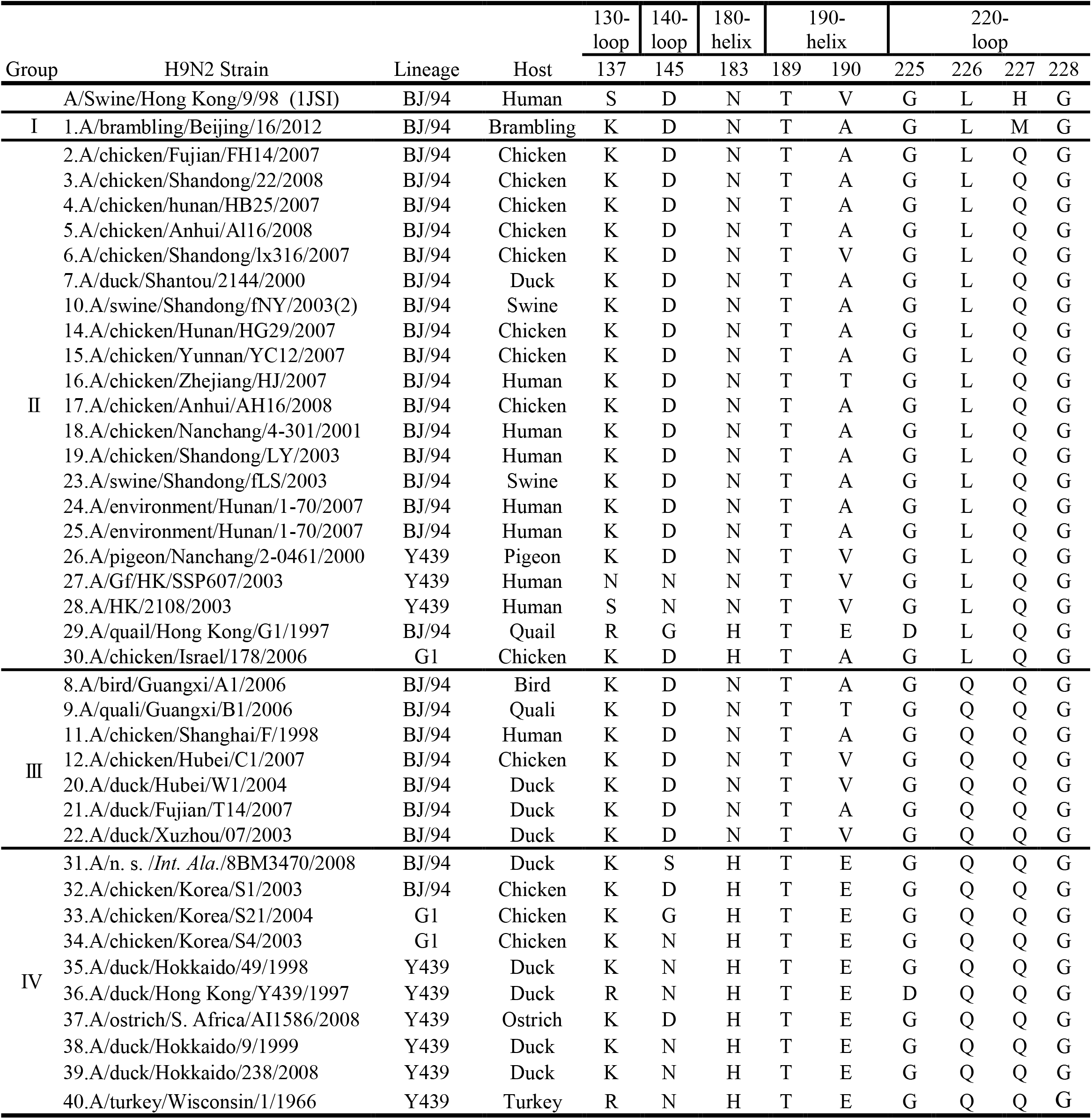
The H9N2 strains are grouped according to the mutations around the binding pocket.

The first group had only one member, A/brambling/Beijing/16/2012. In this model[13], the binding pocket has four highly conservative residues Tyr98, Ser136, Trp153, Leu190while there key mutations locate around LSTc: S137K, H227M and V190A (Figure 2a). In particular, the mutation S137K introduced a salt bridge between the negatively charged sialic acid unit of LSTc and the positively charged side chain of residue Lys137, which significantly stabilized the conformation of sialic acid in the pocket. This mutation also eliminated the hydrogen bond initially established between H227 and R220, which in turn weakened the contact between the two hemicycles of the 220-loop ring, and formed two new hydrogen bonds between the ring and a water molecule previously bound to the ligand. Compared to the hydrogen bond conformation in the X-ray structure (PDB code 1JSI), the water molecule rotated 90° and the bonding atom was shifted from Sia-1-O9 of LSTc to Sia-1-O8. And as a result, 220-loop moves closer to the ligand and establishes stronger interactions with LSTc. Mutation V190A reduces the Van der Waals radius of the side-chain that drive 190-helix close to the GlcNAc-3, however, the introduction of V190A did not bring observable change in the overall structure.

**Figure 2.**
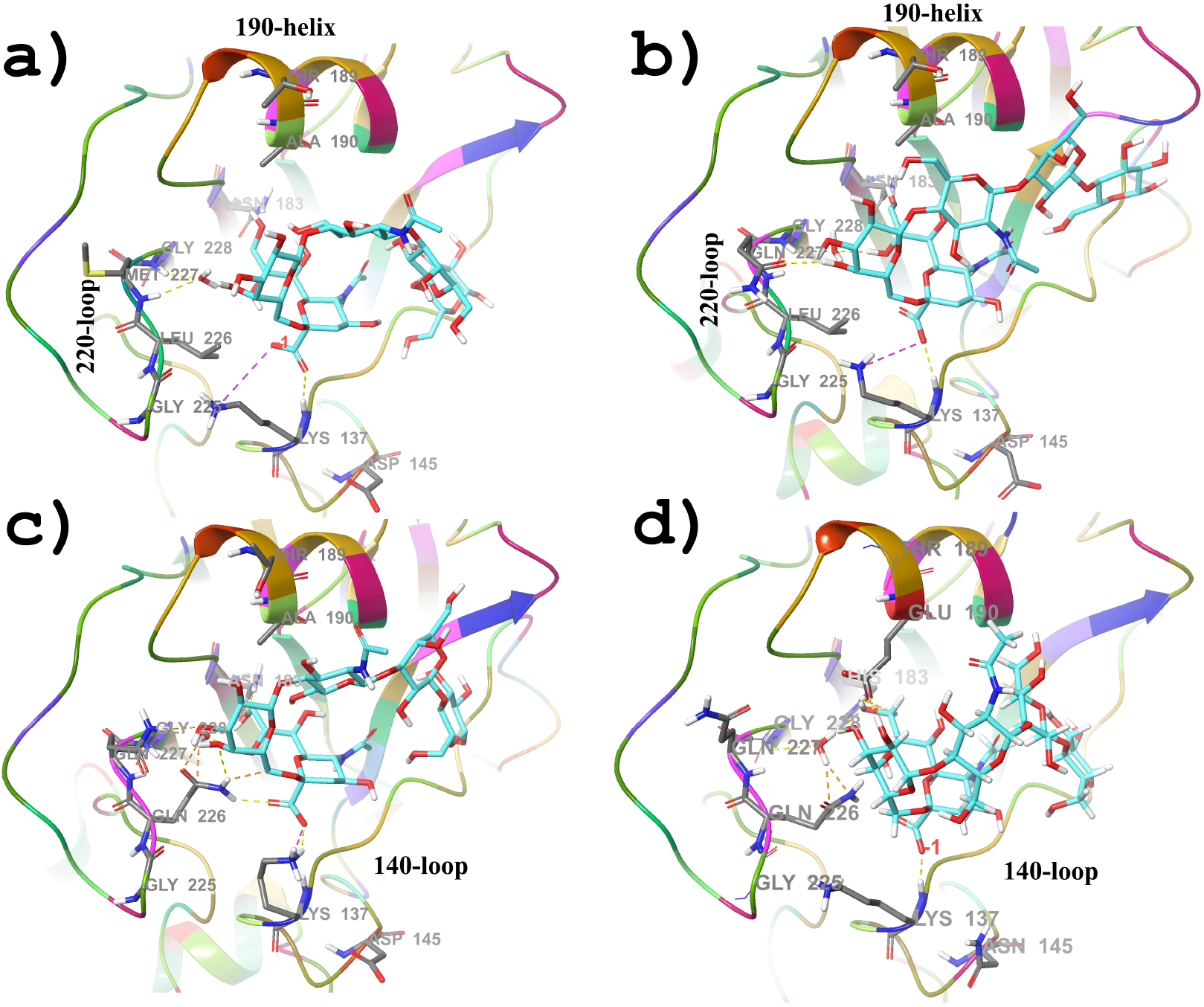
The mutations are categorized into four groups based on the structural characteristics and their effects they have on the HA1-LSTc binding interactions. a: Group I, b: Group II, c: Group III and d: Group IV.

The second group had the largest number of strains studied, which were mainly characterized by the introduction of mutant strains H227Q and S137K, while the key residues L226 remained unchanged. These mutations introduce strong interactions between the amine groups on the side chain and the hydroxyl group of Gal-2, bringing the most twisted region of the ligand (between Gal-2 and Sia-1) closer to 220-loop (Figure 2b). Interestingly, this mutation had little effect on the Sia-1 conformation in the binding pocket and had very limited perturbation to the interaction between the key residue L226 and the ligand. This group also included strains with a V190A mutation that did not cause significant perturbation of the ligand conformation, as observed in group 1.

In the third group, both of the two adjacent residues L226 and H227 were replaced by glutamine, and all strains had the same amino acid distribution at the nine key sites except for some different mutations at site 190 (Figure 2c). Similar to group 2, the second saccharine unit, Gal2, was pulled towards Q227 in most members of the group 3. Compared with the X-ray structure, mutation L226Q made the interaction between side-chain of residue at 226 and Sia-1 more intense. In particular, in the case of A/duck/Fujian/T14/2007, Sia-1 was even pushed to the edge of the binding pocket, where Sia-1 was observed deep inside the bonding pocket in most other model structures, including the X-ray crystal structure. Taken together, the double mutation of L226Q and H227Q introduce relatively large conformation changes to the ligand in the binding pocket. Since all members of this group had nearly identical sequences at all nine key sites, the conformational instability of the LSTc can be attributed to mutations outside the binding pocket, especially the resides at 226 and 227.

All the remaining H9N2 strains studied formed the fourth group, which had two key mutations compared to the three above: A/V190E and N183H. Calculations showed that mutation A/V190E enables the residual 190 to form various contacts with the ligand, building hydrogen bonds between E190-OE and Sia-1-O7/O9, Gal-2-O2/O3/O4, GlcNac-3-O3. These contacts made the ligand-binding conformation more complex than in the third group. The mutation N183H introduced the big imidazole side chain, which in many cases pushed Sia-1 towards residue 226, making it easier for Q226 to come into contact with Sia-1 (Figure 2d).

### The geometric characteristics of the binding of residue 226 to LSTc

Residue 226 of HA1 protein has long been considered as key to the recognition of particular types of host cells by influenza virus. For example, Wan and colleagues reported that H9N2 could be cultured in human respiratory epithelial cells [34]. Leu226 of the HA protein were found more infectious than Gln226, since the strains with former HA can grow at a faster rate, up to 100 times the concentration of the latter. Calculations revealed that Q226 and L226 formed similar hydrogen bonding and van der Waals interactions with Sia-1 atoms. However, three heavy atoms of the Leu226 side chain, CD2, CD1 and CG, were all found within 5Å of Sia-1 atoms, while only NE2 atoms of Q226 found within this range. The average contact distance between LSTc and L226 was found less than that between LSTc and Q226, indicating Sia-1 tends to form stronger contact with Leu226 than with Q226.

To further understand the differences between the Q226 and L226 in controlling the adaptability of H9N2 influenza virus to different hosts, we superposed all the binding configurations of LSTc binding to different HA1 proteins. Figure 3 compared the distribution of the Q226-contacted Sia-1 atoms in the local Q226 coordinates with that of the L226-contacted Sia-1 atoms in L226 coordinates. Sia-1 atoms in contact with L226 were found concentrated in a small region in the L226 coordinates. Their orientation relative to L226 were fairly conservative and the span of most of their Z-components was less than 3.2Å. In contrast, the distribution of Sia-1 atoms in contact with Q226 were very divergent and random, the span of their Z-components was 6.5Å. Therefore, compared with Q226, the structural symmetry of L226 side-chain may help to form a conservative binding configuration between Sia-1 and L226-type HA1, and this repeatable pattern enhancing the probability of their binding.

**Figure 3.**
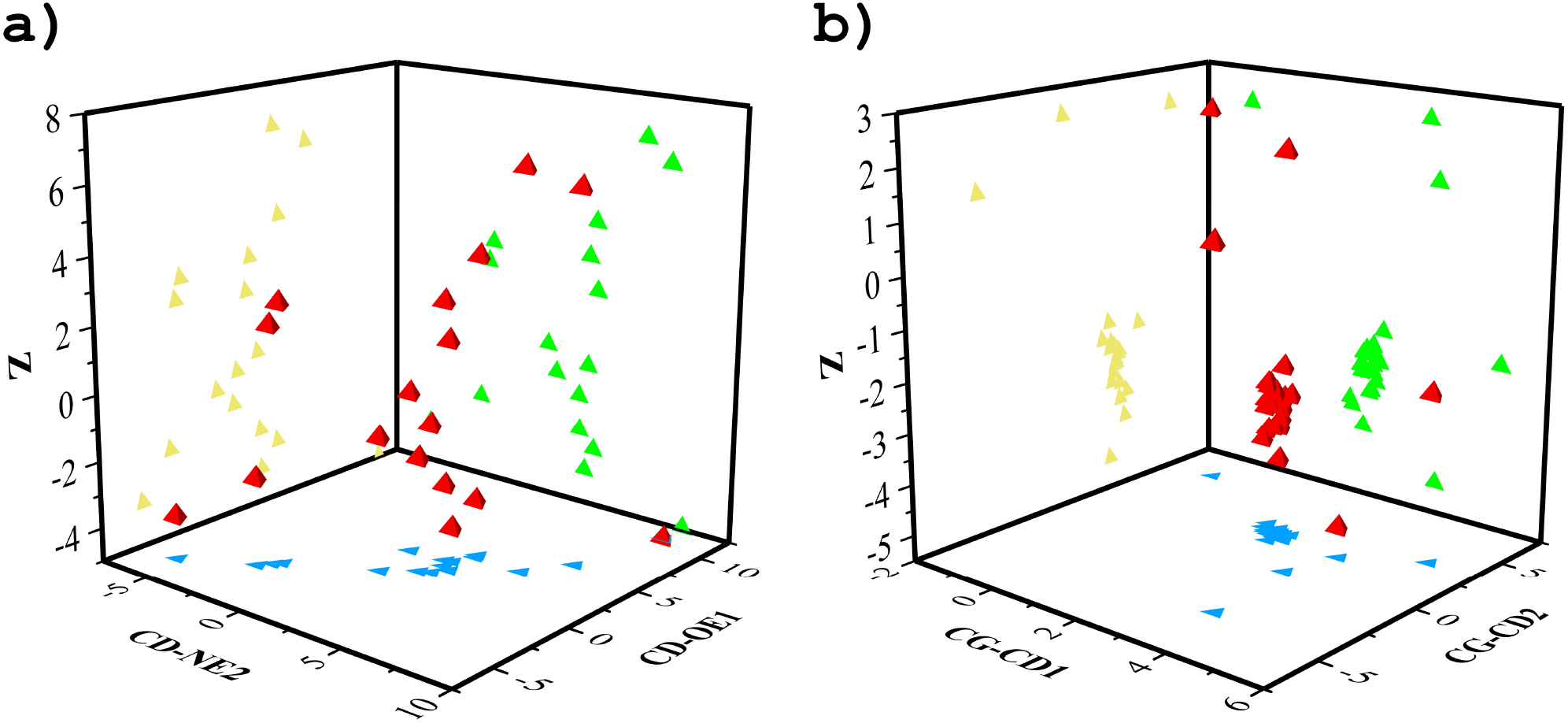
The statistical distribution of the orientation between residue 226 and LSTc is dependent on the mutation. The orientation of the hydroxyl groups hanging outside the Sia-1 sugar ring is: a) variable with respect to the side chain of Q226 and b) relatively stable with respect to the side chain of L226.

### Phylogeny tree built based on the mutations of the key residues

We noted that the HA1 protein of the H9N2 strain under study contained a large number of mutations, with a variety of mutations occurring in nearly 50% of the residues, far beyond the nine mutation sites carefully examined in the above grouping study (Figure S1). Given that these nine mutations are located around LSTc-binding pocket, we wondered whether the sequence of the nine residues largely determined the ability of the virus to infect. To address this problem, we compared the phylogenetic tree built by the sequence of the nine mutations and that constructed from the sequence of the entire H9N2-HA1 protein (Figure 4). First, all of the members of group IV, characterized by two mutations of A/V190E and N183H, were well separated from those of the other groups in both phylogenetic trees. The only exception was A/quail/Hong Kong/ g1/1997 in the second group, which was assigned to group IV in key-residue-based phylogenetic tree of Figure 4a. As a comparison, this strain was correctly separated from the members of group IV but still located closest to group IV in the whole-sequence-based phylogenetic tree of Figure 4b. Second, the strain A/brambling/Beijing/16/2012 of group I stayed closely with the same subset of Group II members. Third, in both phylogenetic trees, the relationship between Group III and II was very complicated, and members from these two groups combined to form a subtree. Moreover, in both subtrees, the members of Group III were divided similarly into two groups, with one group at the upper boundary of the subtree and the other at the lower boundary. Finally, in both trees, the deepest branches were composed of only the members of group II, most of which were strains collected from 2007 and 2008. Taken together, the two types of trees shared very similar topological structure, indicating that the 9 key sites are important not only in binding the ligand but also in the evolution processing.

**Figure 4.**
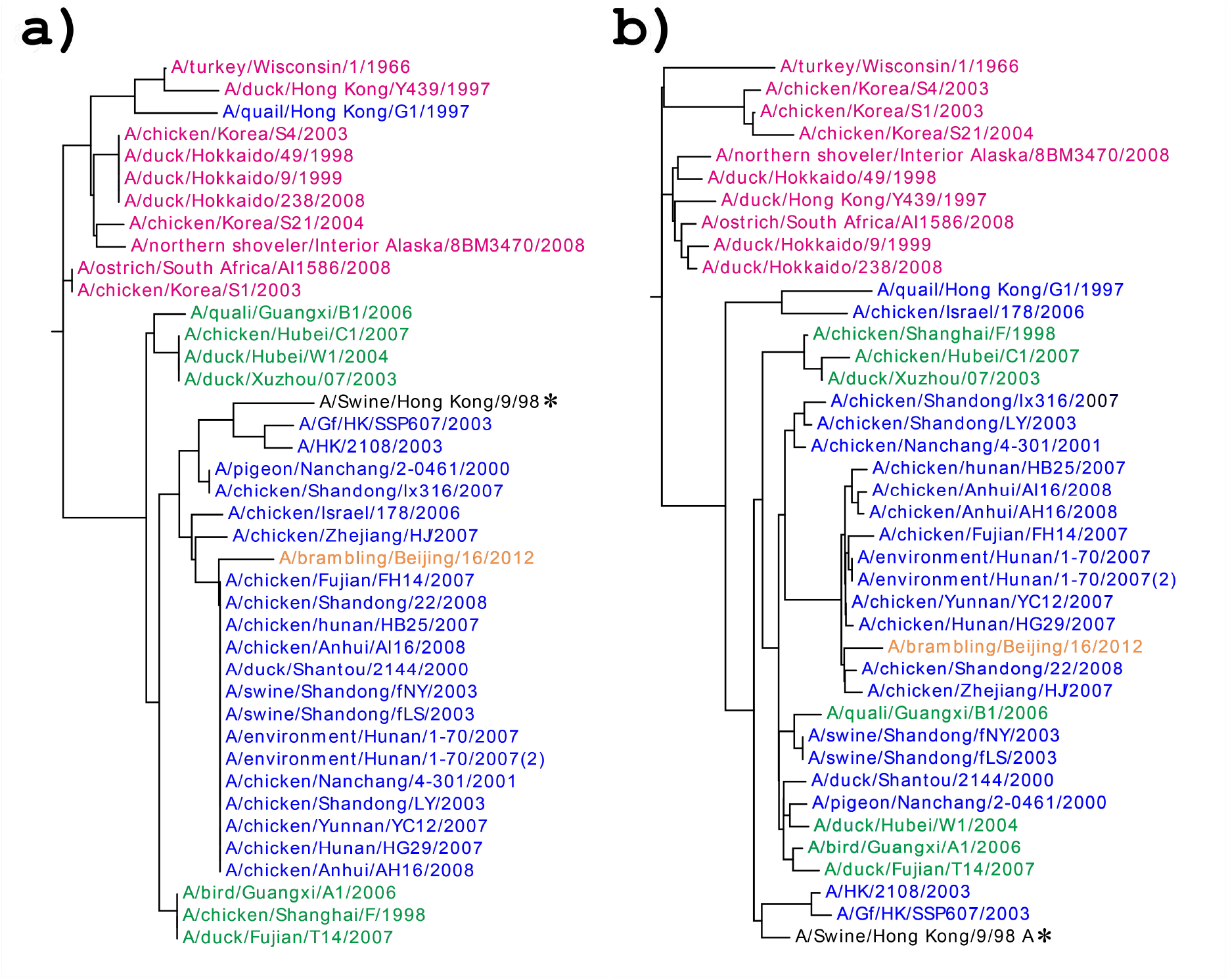
Comparison of phylogeny tree of the studied H9N2 strains. The phylogenetic tree is: a) built using the amino-acid sequences at the nine mutation-site involving LSTc binding, b) using the whole sequence HA1 proteins. The strains are colored as following: brown for group I, blue for group II, green for group III, red for group IV.

## Conclusion

In this study, we compared 40 HA protein structures of AIV H9N2 virus, mostly found in China, to study their ability to bind to LSTc as a first step in infecting mammals. The three-dimensional structures of HA1 proteins of 40 H9N2 virus were built and their interactions with LSTc were analyzed via docking simulation, in particular by characterizing key mutations around the ligand-binding pocket. We categorized the mutations according to the perturbations they produced, which are mainly divided into those involved in direct ligand-binding interaction and those involved in indirect interaction. Both 200-loop and 190-Helix are intermediate elements that regulate the effect of mutations on LSTc-binding. The importance of the selected mutations was demonstrated by the fact that the phylogenetic tree constructed from the selected mutant sequences had a similar topological structure to that made from the entire protein sequence. Our calculations suggested that examining the molecular microenvironment perturbed by key mutations might provide a way to understand how the virus acquires the ability to infect humans through mutation at the molecular level.

## Acknowledgements

This work was supported, in part, by the National Key Research and Development Program of China (Grant No. 2017YFC1600900, 2019YFA0905701) and by the Key University Science Research Project of Jiangsu Province (Grant No. 17KJA180005). X.H. was supported by National Students’ platform for innovation and entrepreneurship training program (Grant No. 201910291055Z).

## Supporting information

Table S1. list of the studied H9N2 strains

Figure S1. The alignment of the selected 40 H9N2 influenza A virus strains.

